# An Axon-Pathfinding Mechanism Preserves Epithelial Tissue Integrity

**DOI:** 10.1101/2020.04.29.068387

**Authors:** Christian M. Cammarota, Tara M. Finegan, Tyler J. Wilson, Sifan Yang, Dan T. Bergstralh

**Author notes:** These authors contributed equally (Christian M. Cammarota, Tara M. Finegan).

## Abstract

Dividing cells often move apically within epithelial tissue layers, likely to escape the spatial confinement of their neighbors. Because of this movement, daughter cells may be born displaced from the tissue layer. Reintegration of these displaced cells helps support tissue growth and maintain tissue architecture. In the *Drosophila* follicular epithelium, reintegration relies on the immunoglobulin-superfamily cell-adhesion molecules (IgCAMs) Neuroglian and Fasciclin 2, which line cell-cell borders^1^. These molecules have been described in epithelia, but are well-studied for their roles in neural development^2–8^. We show here that reintegration works in the same way as IgCAM-mediated axon growth and pathfinding; it relies not only on extracellular adhesion but also mechanical coupling between IgCAMs and the lateral Spectrin-Based Membrane Skeleton. Our work indicates that reintegration is mediated by a distinct epithelial cell-cell junction that is compositionally and functionally equivalent to junctions made between axons.

## Introduction

Epithelial tissues form the boundaries of organs, where they perform a range of functions, including secretion, absorption, and protection. These tissues are commonly comprised of discrete cell layers - sheets of cells that are one-cell thick. Cell proliferation, a fundamental process in epithelial tissue homeostasis and remodeling, poses a potential challenge to the integrity of these layers; the division process, which features cell growth, cell shape change (rounding), and cytokinesis, occurs within an already tightly-packed sheet. In multiple systems examined, dividing epithelial cells round up and change their apical-basal (z-axis) position within the tissue. This movement is likely a response to neighbor-cell crowding, and offers the mitotic cell space in which to divide.

Apical-directed movement of the cell nucleus prior to division, called interkinetic nuclear migration (INM), is primarily studied in pseudostratified vertebrate neural tissues, where it is thought to play a role in neural differentiation^9,10^. Similar movement is also evident in columnar epithelia, including the lining of the mammalian small intestine^11,12^. A striking example occurs in the ureteric bud. In that tissue, mitotic epithelial cells apparently leave the layer, moving into the lumen to divide ^13^. Protrusion of mitotic cells is also observed in the cuboidal *Drosophila* follicular epithelium (FE), and a daughter cell can be born mostly outside the layer^1^. As in the ureteric bud, the misplaced cell is not lost, but rather reintegrates into the tissue.

Two conserved IgCAMs, Neuroglian (Nrg) and Fasciclin 2 (Fas2), participate in cell reintegration in the *Drosophila* follicular epithelium^14^. Nrg and Fas2 are enriched along follicle cell-cell borders during the developmental time over which follicle cells are dividing, with expression dropping off soon afterward^15,16^. When these factors are disrupted, cells can fail to reintegrate and instead remain positioned above (apical to) the epithelial layer (Figure 1A). We refer to this phenotype as “popping out.”

**Figure 1.**
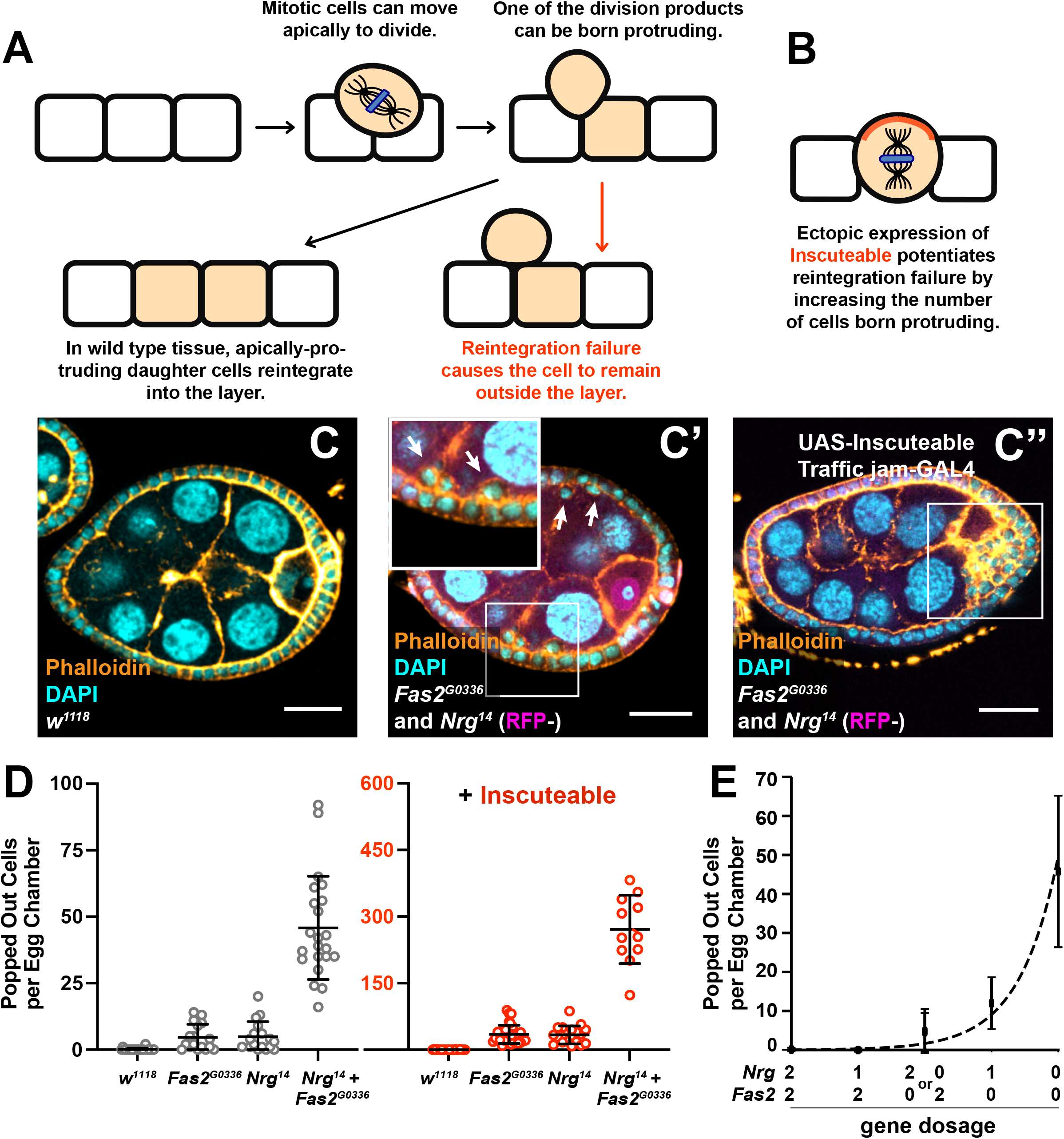
Nrg and Fas2 act in parallel to drive reintegration. **A)** Diagram showing apical cell movement during mitosis and subsequent cell reintegration. **B)** Inscuteable increases out-of-plane divisions, and therefore the number of reintegration events. **C and D)** Genetic disruption of both *Fas2* and *Nrg* causes a synergistic increase in reintegration failure over disruption of either gene alone. Ectopic expression of Inscuteable has a multiplicative effect on reintegration failure. Representative images are shown in C, quantification in D. Each data point represents the number of extralayer (popped out) cells in one egg chamber. Each column represents three or more dissections of five or more flies. Significance was determined using an unpaired, two-tailed student’s t-test with Welch’s correction. Scale bars in B correspond to 20 μm. **E)** The relationship between reintegration failure and gene function is not linear. A plot of gene dosage *versus* misplaced cell number fits an exponential curve (R^2^ = 0.972), providing a speculative model for reintegration function.

Like their vertebrate orthologs L1CAM and NCAM1/2 (respectively), Nrg and Fas2 are cell adhesion molecules primarily studied in the context of nervous system development, in which they promote neurite fasciculation, synaptogenesis, and axon guidance^2–5^. Both proteins have also been identified as components of the insect septate junction, which is the functional equivalent of the vertebrate tight junction^17^. However, proliferating follicle cells do not have mature septate junctions, which suggests that Nrg and Fas2 participate in a distinct adhesion assembly in the FE^18^. In this study we set out to explore the composition and function of this assembly. Our findings indicate that it is equivalent in both aspects to the axon-axon junctions that drive axon growth.

### Fas2 and Nrg work in parallel to mediate reintegration

Nrg and Fas2 both mediate reintegration in the FE, but whether they act in parallel or interdependently has not been tested^14^. We disrupted them genetically using the null alleles *Nrg*^*14*^ or *Fas2*^*G0336*^ and measured the number of popped-out cells in Stage 6-8 egg chambers. Whereas genetic loss of either Fas2 or Nrg causes ~5 popped-out cells per egg chamber (~1/150 cells), losing both causes ~50 popped-out cells per egg chamber (~1/15 cells) (Figure 1C,D and Supplemental Figure 1).

We tested whether this effect reflected an increase in failed reintegration events. Ectopic expression of Inscuteable (Insc) potentiates reintegration failure because Insc reorients mitotic spindles in the FE, increasing the frequency with which a daughter cell is born apical to the tissue layer (Figure 1B)^14^. Inscuteable expression has a multiplicative effect on popping out, increasing the number of misplaced cells in tissue mutant for either *Nrg*^*14*^ or *Fas2*^*G0336*^ by ≅ six-fold (Figure 1D). Because Inscuteable has the same effect for *Nrg*^*14*^, *Fas2*^*G0336*^ double mutants, we conclude that the increase in popped-out cells in double mutant tissue represents an increase in failed reintegrations, and therefore that Nrg and Fas2 act in parallel (Figure 1C,D). These proteins are likewise functionally redundant in the control of axon guidance^19^.

While *Nrg* and *Fas2* mutants have identical phenotypes, indicating equal contribution to reintegration, the loss of both factors causes a synergistic rather than additive increase in reintegration failure. To investigate the relationship between reintegration function and Fas2/Nrg activity, we built a speculative mathematical model by plotting popped-out cells versus gene dosage on a Cartesian plane. The assumption that function correlates with gene dosage has not been tested directly, but agrees with the correlation between *Nrg* dosage and Nrg expression measured in Figure S2. To expand our data set, we used a genomic-rescue strategy to restore one copy of *Nrg* in *Nrg*^*14*^ single and *Nrg*^*14*^, *Fas2*^*G0336*^ double mutants. The resulting data points fit an exponential curve (Figure 1E). This curve suggests that reintegration is robust, as it is typically successful even at low IgCAM function.

### The cytoplasmic FIGQY subsequence of Neuroglian participates in reintegration

*Nrg*^*14*^ and *Fas2*^*G0336*^ have indistinguishable phenotypes (Figure 1), which raises the question of whether the proteins have different functions in mediating reintegration. We explored this question by using a genomic-rescue strategy to express Nrg in *Fas2*^*G0336*^ and *Nrg*^*14*^ mutant backgrounds^20^. Ectopic expression of Insc was used to sensitize the popping out phenotype. Expression of Nrg in *Nrg*^*14*^ tissue rescues reintegration failure completely, regardless of Insc expression (Figure 2A). Expression of Nrg in *Fas2*^*G0336*^ tissue rescues reintegration, but sensitization with Insc shows that the restoration of function is incomplete. These results are consistent with the possibility that Nrg and Fas2 have both overlapping and separate functions.

**Figure 2.**
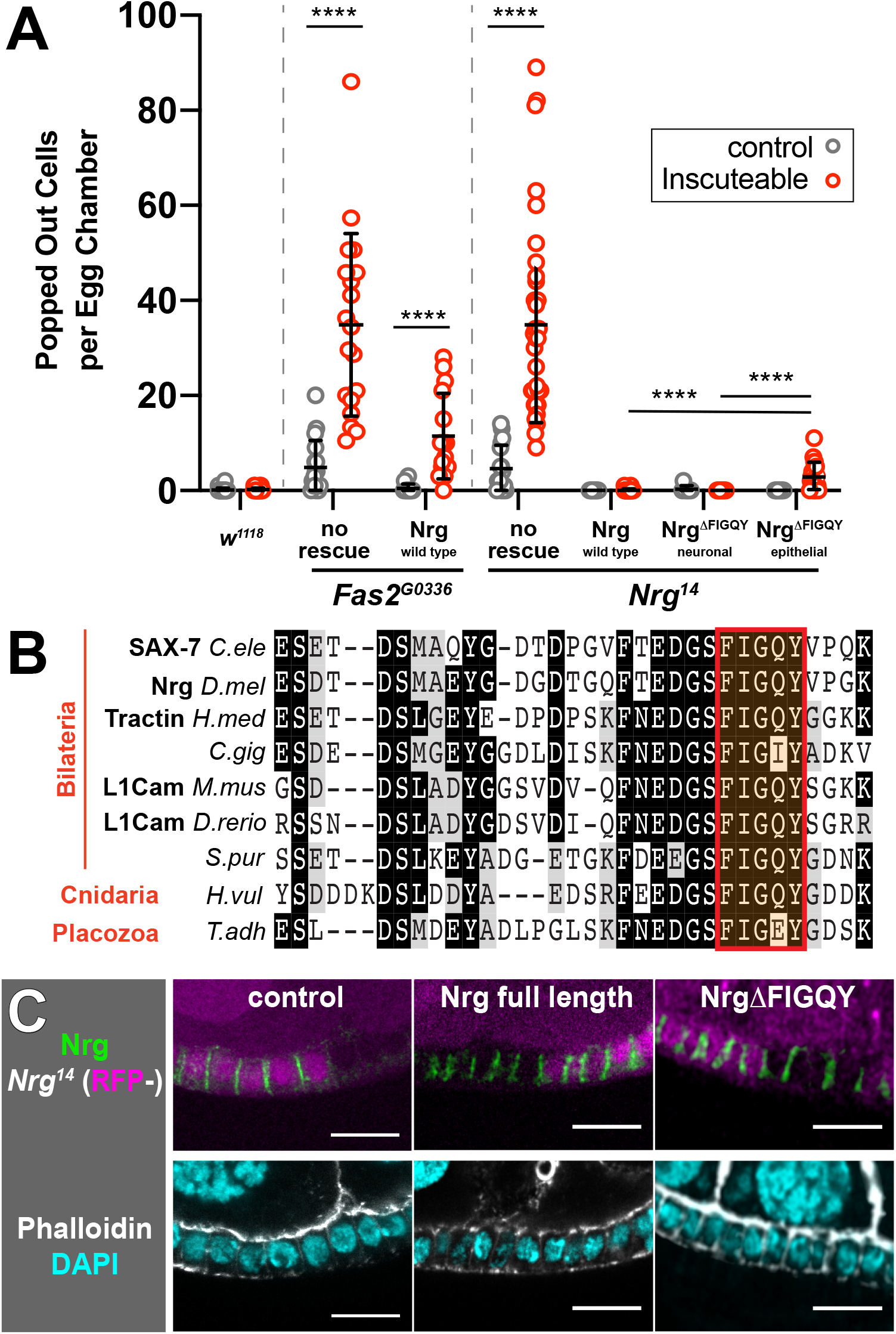
The cytoplasmic FIGQY sequence of Nrg participates in reintegration. **A)** Neuroglian rescues reintegration failure in *Nrg* null tissue, but only partially rescues reintegration failure in *Fas2* null tissue. Removal of the FIGQY subsequence from the epithelial Nrg isoform allows for partial rescue in *Nrg* null tissue. Neuroglian and Nrg variants were expressed from a single genomic rescue construct. The neuronal isoform (180) includes an alternate exon that encodes the FIGQY subsequence, and the epithelial isoform (167) should not be affected. Quantification and statistical tests performed as in Figure 1. **B)** The Ankyrin-binding domain of Nrg/L1CAM is evolutionarily conserved. **C)** The FIGQY subsequence is not required for Nrg localization to epithelial cell-cell borders. Representative images are shown in C, and quantified in Supplemental Figure 2. Scale bars correspond to 5 μm.

Since both Nrg and Fas2 are cell-cell adhesion proteins, we considered whether the basis for their distinction was intracellular. The cytoplasmic C-terminus of Nrg/L1CAM includes a highly conserved five amino acid subsequence (FIGQY) that helps mediate interaction with Ankyrin, a linker between the cytoskeleton and the membrane^21,22^. Deletion of FIGQY reduces *in vitro* interaction between Ankyrin and the vertebrate L1CAM paralog Neurofascin by 80%^22^. The remaining contact requires FIGQY-adjacent amino acids, which are also highly conserved (Figure 2B)^22^.

We tested whether FIGQY contributes to reintegration by expressing Nrg^ΔFIGQY^ transgenes in an *Nrg*^*14*^ background^20^. In both insects and vertebrates, one Nrg/L1CAM isoform is expressed in epithelia (*Drosophila* Nrg167) and a different isoform in neural tissue (Nrg180)^23^. Reintegration failure is completely rescued if FIGQY is deleted from only the Nrg180 isoform (leaving Nrg167 intact) (Figure 2A). Deletion of FIGQY from only the Nrg167 isoform partially rescues reintegration failure in the presence of Insc. Our model suggests that this transgene provides only ~35% of wild-type function (Figures 1C and 2A).

We considered whether removal of the FIGQY subsequence affects localization of Nrg to cell-cell contacts. Nrg167^ΔFIGQY^ localizes to follicle cell-cell borders in *Nrg*^*14*^ mutant clones, as revealed by immunostaining (Figure 2C). Quantification of these images shows that the relative strength of Nrg immunoreactivity corresponds to genetic dosage; signal decreases by ~25% at the border between a cell expressing two copies of Nrg and a cell expressing only one copy, and ~50% at the border between cells expressing one copy each (Supplemental Figure 2). Notably, Nrg does not require homotypic adhesion to localize to the membrane, as the signal decreases by only half at the border between wild-type and *Nrg*^*14*^ clones. These results are consistent with observations made in other organisms; neither the cytoplasmic domain of L1CAM nor its *C. elegans* ortholog SAX-7 are required for cell-cell adhesion or membrane localization^24,25^. Together, our findings indicate that the FIGQY sequence mediates an intracellular function involved in cell reintegration.

### Neuroglian mediates reintegration through interaction with Ankyrin

During axon growth, Ankyrin stabilizes Nrg/L1CAM at the cell surface by linking it to the Spectrin-Based Membrane Skeleton (SBMS), a subcortical scaffold that mechanically supports the plasma membrane^20^. Ankyrin therefore acts as a molecular ‘clutch’ by which new outgrowth is mechanically stabilized^28^. We tested whether this mechanism also participates in cell reintegration.

Like Neuroglian and Fas2, Ankyrin localizes along the length of lateral cell-cell junctions in the FE^26^. We used short-hairpin RNA to knockdown Ankyrin. Though we cannot determine the extent of knockdown directly, this shRNA has been effective in a previous study^27^. Ankyrin knockdown in the FE results in popped-out cells, making it the first non-adhesion protein to give this phenotype (Figure 3A,B). This result is not explained by a decrease in the junctional localization of Nrg and Fas2, as both are unchanged in Ankyrin-shRNA tissue (Supplemental Figures 3A,B). We also find that Ankyrin knockdown potentiates reintegration failure in *Fas2*^*G0336*^, but not *Nrg*^*14*^, tissue (Figure 3B). Together, these findings show that Nrg and Ankyrin work together in parallel to mediate reintegration.

**Figure 3.**
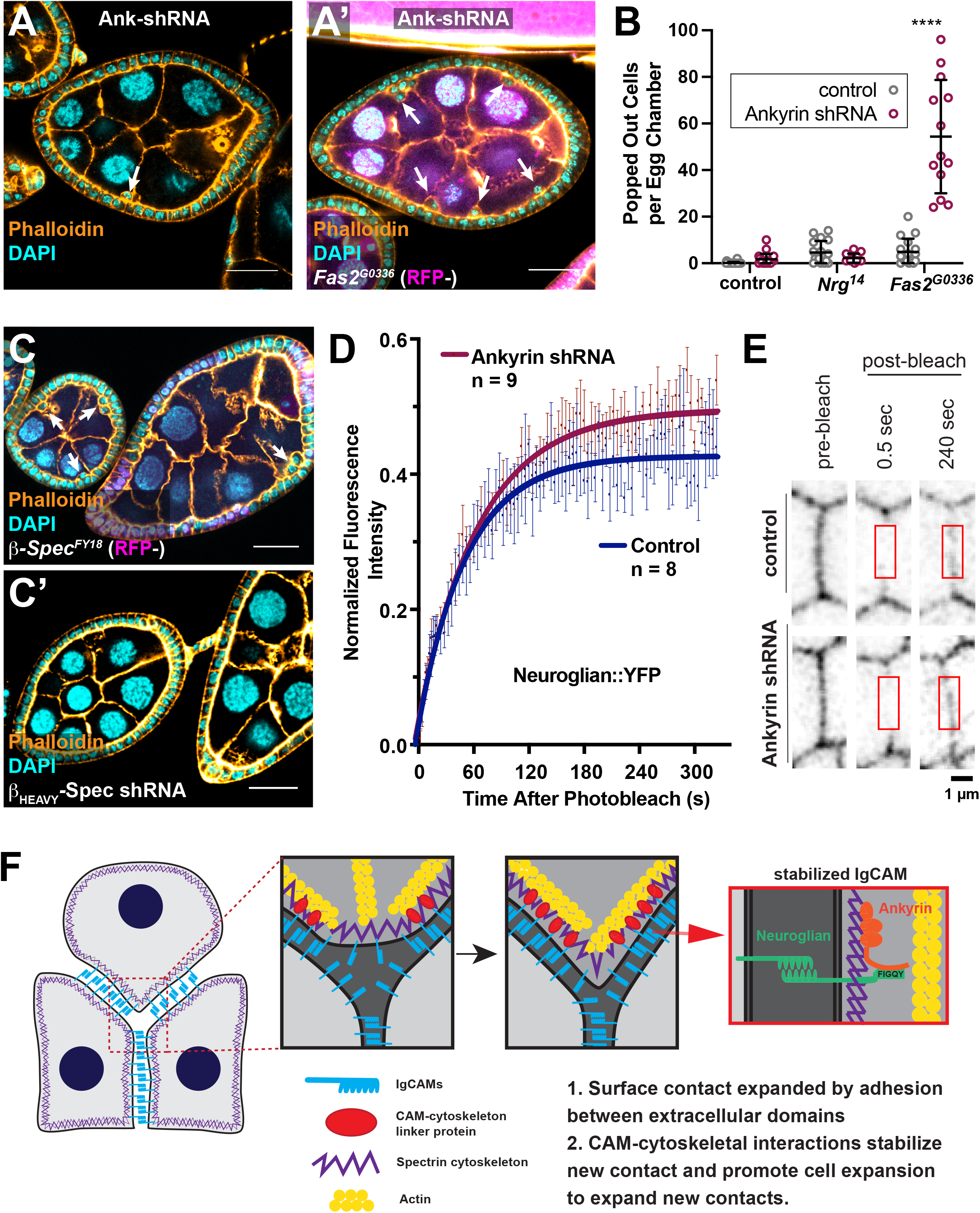
Nrg and Ankyrin cooperate to drive cell reintegration. **A and B)** Genetic disruption of Ankyrin results in failed cell reintegration. Expression of Ankyrin shRNA, driven by Traffic Jam-GAL4, results in ~2 popped out cells per egg chamber. Ankyrin shRNA potentiates the reintegration failure phenotype seen in *Fas2* null tissue, but not *Nrg* null tissue. Representative images are shown in A, quantification in B. Quantification and statistical tests performed as in Figure 1. Scale bars in A correspond to 20 μm. **C)** Popped-out cells, indicating reintegration failure, are apparent in FE tissue mutant for β-Spectrin^FY18^ but not β_heavy_-Spectrin. Scale bars correspond to 20 μm. **D)** Fluorescence recovery curves of Nrg::YFP after photobleaching. Lateral FE cell junctions were bleached in egg chambers expressing Ankyrin-shRNA driven by Traffic Jam-GAL4 or egg chambers with the driver alone (control). Fluorescence intensity was normalized to take into account inherent photobleaching due to imaging. Experiments were performed independently with number of experimental repeats indicated. Curves were calculated by fitting a one phase association curve. Error bars represent SEM. The 95% confidence interval (CI) of the control plateau is 41-44. This does not overlap with the 95% CI of the control plateau, which is 48-51. **E)** Frames from an example FRAP experiment, showing the fluorescence of Nrg::YFP at a cell junction prior to and post bleaching in control and Ankyrin-shRNA expressing egg chambers. Red boxes indicate photobleached region over which fluorescence intensity was quantified. **F)** We propose a refined model for Ig-CAM mediated cell reintegration in epithelia, inspired by existing knowledge of how axon growth occurs. Both extracellular adhesion and intercellular connection to the cytoskeleton is necessary for the reintegration of cells into epithelial layers. Specifically, we reveal that the spectrin-based cytoskeleton stabilizes Nrg cell-cell adhesion through Ankyrin to facilitate reintegration. We propose that the SBMS mechanically stabilizes the trans-interactions of Nrg at the ‘leading edge’ of integrating cells to facilitate the progression of cell reintegration. Nrg-Ankyrin interactions thereby provide a traction force (grip).

The SBMS, a lattice comprised of ◻- and β-spectrin tetramers, is polarized in epithelial cells. In the FE, β-spectrin (called β-Fodrin or βII-spectrin in vertebrates) and Ankyrin line the lateral cortex, whereas β_heavy_-spectrin is along the cell apex, from which Ankyrin is excluded^26,29–32^. Several studies demonstrate disorganization in *β-spectrin* mutant follicular epithelia^33–35^. In agreement, we observed extensive disorganization phenotypes in large mitotic *β-spectrin*^*FY18*^, particularly after Stage 6 of egg chamber development (not shown). However, we also observed popped-out cells in earlier, smaller clones (Figure 3C). Although we cannot exclude the possibility that this phenotype relates to reported defects in Hippo signaling or actomyosin that cause later disorganization, it is consistent with a failure of cell reintegration. The localization of Nrg and Fas2 to cell-cell borders is unchanged in *β-spectrin*^*FY18*^ mutant clones (Supplemental Figure 3C,D). Popped-out cells were not observed after knockdown of β_heavy_-spectrin using an shRNA that has been effective in a previous study^36^ (Figure 3C).

To test whether Ankyrin affects Nrg mobility in epithelia, we used Fluorescence Recovery After Photobleaching (FRAP) analysis to measure the effect of Ankyrin disruption on Nrg stability at follicle cell-cell junctions. In the control condition, Nrg::YFP signal recovers to only 42% of its pre-bleach intensity, indicating that the majority of Nrg is immobile (Figure 3D,E). Ankyrin-knockdown increases signal recovery to 50%, indicating that Ankyrin-binding contributes to some, though not all, of the immobile fraction (Figure 3D,E). These findings are consistent with observations made in motoneurons, in which deletion of the FIGQY sequence increases the mobility of Nrg^20^. For technical reasons, our experiment was performed at interphase cell-cell junctions, leaving open the possibility that Ankyrin has an even stronger effect in actively reintegrating cells.

Together, these results are consistent with a model in which Nrg cooperates with the lateral SBMS to provide a traction force (grip) during cell reintegration, as it does during axon growth (Figure 3F).

### The intracellular region of Fas2 participates in reintegration

We next tested whether Fas2, like Nrg, has an intracellular function in reintegration. There are seven isoforms of Fas2 in *Drosophila.* Five of these have transmembrane domains, and are considered to be neuronal^37^. The other two, which have been studied in non-neuronal cells in the trachea, Malpighian (renal) tubule, and in glial cells, are tethered to the membrane by glycosylphosphatidylinositol (GPI) anchors, indicating that they have only adhesive function^5,37–39^. Since the FE is not neuronal, these observations suggest the possibility that Fas2 might mediate epithelial cell reintegration through adhesion alone. However, the FE expresses at least one “neuronal” isoform (*i.e.*, including a transmembrane domain), as revealed by previous immunostaining with the 1D4 monoclonal antibody^1,2,37^.

We tested whether the cytoplasmic domain of Fas2 is important for epithelial cell reintegration by expressing YFP-tagged UAS-Fas2 variants in the FE. UAS-Fas2-Full-Length, which includes a cytoplasmic region (isoform PD), rescues reintegration failure in *Nrg*^*14*^ and *Fas2*^*G0336*^ tissue, consistent with the finding that Nrg rescues both these mutants (Figures 2A and 4A). UAS-Fas2-Extracellular, which includes the extracellular and transmembrane regions but lacks an intracellular domain, rescues a non-neuronal function of Fas2, namely the regulation of brush border length in the Malpighian tubule epithelium^38^. However, we found that neither UAS-Fas2-Extracellular nor UAS-Fas2-Intracellular (the transmembrane and intracellular region of isoform PD) rescue reintegration failure in *Fas2*^*G0336*^ tissue, though both variants localize at lateral cell junctions (Fig 4A,B). These results show that the reintegration role played by Fas2 requires a combination of its adhesive and cytoplasmic functions.

**Figure 4.**
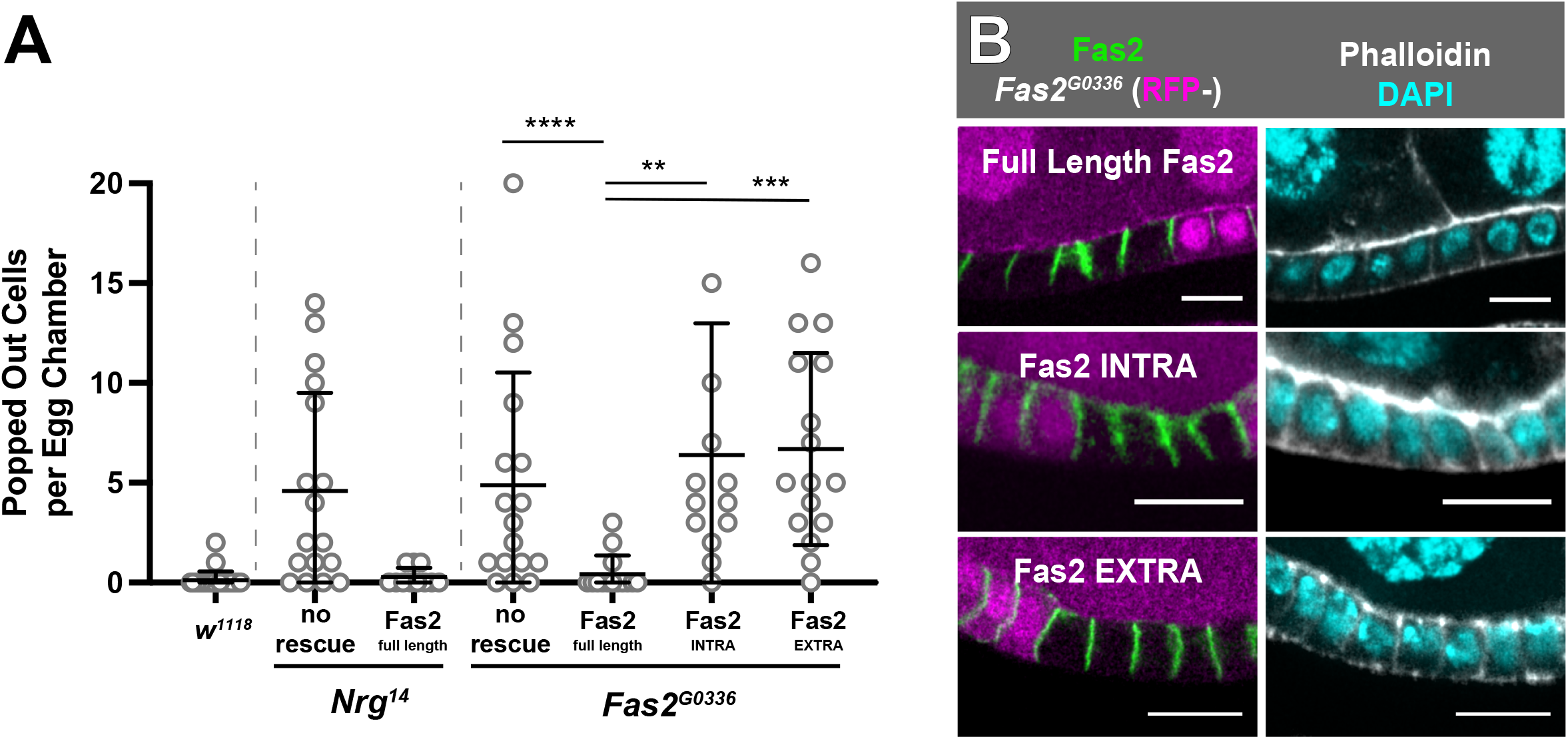
Both the extracellular and intracellular domains of Fas2 participate in reintegration. **A)** Expression of full length Fas2 rescues reintegration failure in *Nrg* and *Fas2* null tissue, but both the extracellular and intracellular domains are required to rescue *Fas2*^*G0336*^. UAS-Fas2-YFP and variants were expressed using the driver Traffic Jam-GAL4. Quantification and statistical tests performed as in Figure 1. The question of whether Fas2 rescues *Nrg*^*14*^ in the presence of Inscuteable was not addressed, as we could not be convinced that the pool of GAL4 was sufficient to drive full expression of both UAS-Fas2 and UAS-Inscuteable. **B)** Fas2 localizes to cell-cell borders without either its extracellular or intracellular domain. UAS-Fas2-YFP and its truncated variants, expressed using Traffic Jam-Gal4, could all be detected at follicle cell-cell borders, whether in the presence (RFP+) or absence (RFP−) of endogenous Fas2. Scale bars correspond to 5 μm.

### Fas3 is a reintegration factor, but NrxIV is not

Like Nrg and Fas2, Fas3 is a homophilic IgCAM found in neurons and in septate junctions^6–8,40,41^. We observe that Fas3 localizes along epithelial cell-cell junctions in early-stage *Drosophila* egg chambers, but expression drops off after follicle cell division ceases at Stage 6 (Figure 5A). This pattern, which is also seen for Nrg and Fas2, suggests a role for Fas3 in proliferation^1,15,16^.

**Figure 5.**
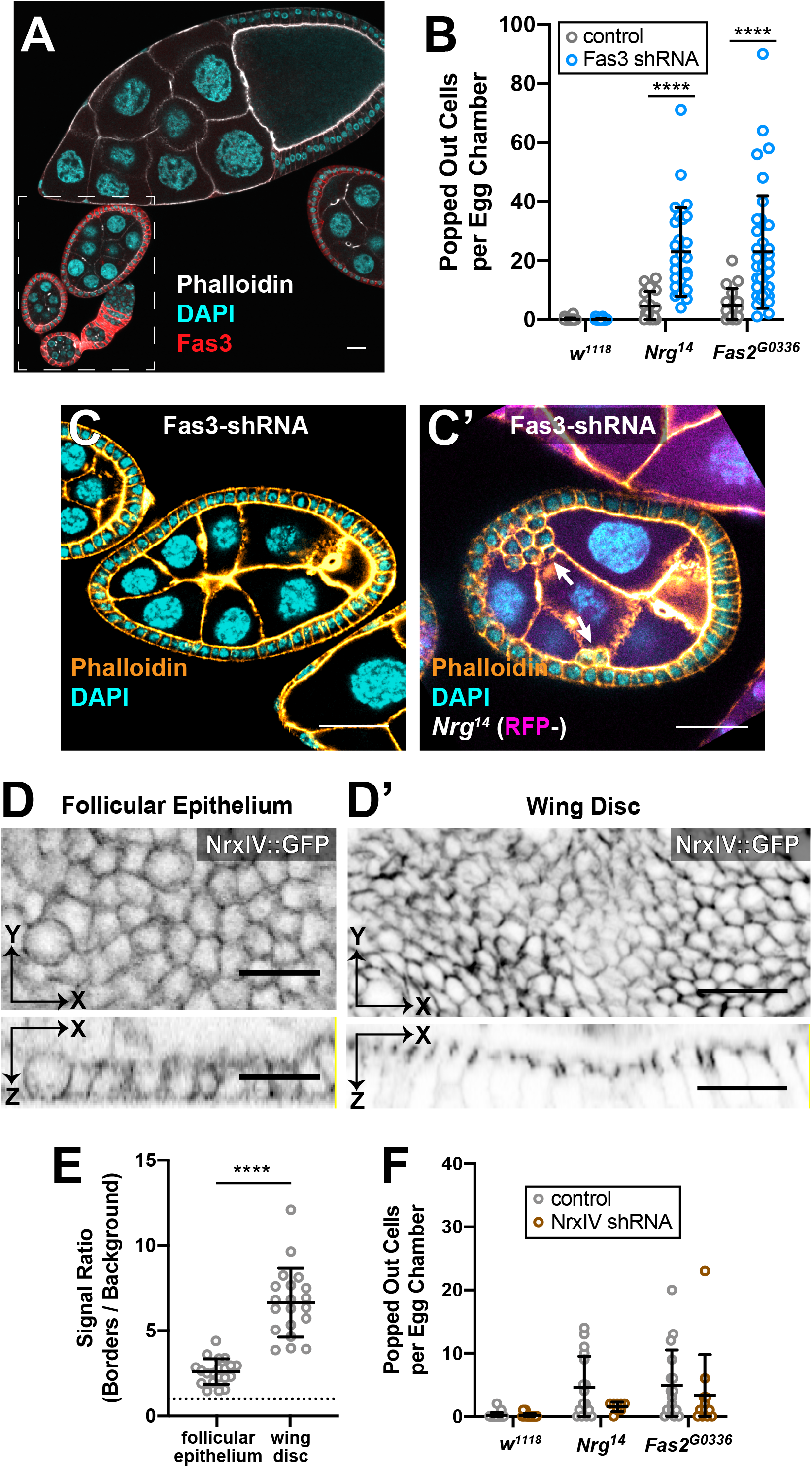
Fas3 participates in reintegration, but NrxIV does not. **A)** Fas3 localizes to follicle cell-cell borders in early stage egg chambers. Fas3 immunoreactivity is detected in early egg chambers, in which the follicular epithelium is proliferative (Stages 1-6, in dashed box), but drops off substantially by mid-stage (Stage 10, top of panel). **B and C)** Fas3 disruption potentiates reintegration failure in *Nrg* and *Fas2* mutant tissue. Fas3 knockdown does not result in misplaced cells (B) unless combined with mutation of either Nrg or Fas2 (B and C). Fas3-shRNA was driven with Traffic Jam-GAL4. Misplaced cells were quantified as in Figure 1. **D and E)** NrxIV is weakly localized in the early follicular epithelium. NrxIV::GFP, a protein trap, demonstrates weaker junctional localization in the follicular epithelium (D) than in the mature wing disc (D’) relative to local background signal (quantification in E). Signal intensity was measured at septate junctions (wing disc) or along lateral cell-cell contacts (follicular epithelium) and compared with adjacent cytoplasmic signal. **F)** NrxIV disruption does not enhance reintegration failure. NrxIV-shRNA, driven with Traffic Jam-GAL4, does not increase the number of misplaced cells in control tissue or in *Nrg* or *Fas2* mutant tissue.

Fas3 has long been implicated in axon guidance, but mutation of *Fas3* alone has no axon guidance phenotype; it is thought to work redundantly with more consequential factors, including Nrg and Fas2^39,40^. Consistent with this, we did not observe popped-out cells in Fas3-shRNA follicular epithelia, but found that Fas3 knockdown increases popping out in *Fas2*^*G0336*^ or *Nrg*^*14*^ mutant tissue by approximately 5-fold (Figure 5B,C). Immunostaining confirmed the efficacy of knockdown (Supplemental Figure 4A). Since Insc has a similar potentiating effect, we tested whether Fas3-shRNA alters spindle orientation in the FE. Spindle orientation is unaffected by Fas3 knockdown (Supplemental Figure 4B,C). Together, these results suggest that Fas3 plays analogous roles in axon guidance and reintegration.

The follicular epithelium does not have mature septate junctions, but the observation that reintegration involves three septate junction components – Nrg, Fas2, and Fas3 - suggests the possibility that reintegration relies on an “immature” septate junction. To test this possibility, we examined the core septate junction molecule Neurexin IV (NrxIV)^17,42,43^. Like Nrg and Fas2, Neurexin-family proteins are found at both synapses and mature septate junctions^17,44^. However, unlike Nrg and Fas2, Neurexins are not implicated in axon guidance. Using both endogenously-tagged NrxIV::GFP and immunostaining, we observed NrxIV along follicle cell-cell borders in early stage egg chambers (Figure 5D,E and data not shown). These results confirm NrxIV expression in the FE, though it is significantly weaker (2.6-fold > background, ± 0.9) than that observed at mature septate junctions in the imaginal wing disc (6.7-fold > background, ± 0.4) (Figure 5D,E). Knockdown of NrxIV in the FE did not result in reintegration failure, either alone or in combination with *Nrg*^*14*^ or *Fas2*^*G0336*^ (Figure 5F). The efficacy of NrxIV-shRNA was confirmed in the wing disc (Supplemental Figure 4D). Taken together, these results show that not all septate junction proteins are reintegration factors, and instead suggest that reintegration relies on a subset of IgCAMs that participate in axon guidance.

Our work shows that epithelial cell reintegration relies on a distinct junctional complex, previously considered an “immature” septate junction, that is neither the classical zonula adherens nor the zonula occludens. These junctions are equivalent in composition and function to the axon-axon junctions that support axon growth and pathfinding. One question that is raised is how the reintegration mechanism evolved. Mature epithelial pleated septate junctions are similar in composition to synaptic junctions, and it has long been proposed that these two types of junction share an evolutionary history^17,45–49^. We therefore suggest that the maturation of septate junctions is akin to synaptogenesis. Both processes preserve IgCAM components of the earlier junction, but feature recruitment of analogous proteins, leading to increased junctional complexity and structural stability. Whether these junctions arose in neural cells is unclear. Both Fas2/NCAM and Nrg/LICAM (Figure 2B) have been identified in Placozoa^50^. These animals have epithelial, but not neural cell types, suggesting the possibility that the epithelial functions of these IgCAMs are ancestral.

## Materials and Methods

### Fly stocks and reagents

The *Drosophila melanogaster* strains and the reagents used in this study are described in the Supplemental Material.

### *Drosophila* genetics

Follicle cell clones of *Nrg*^*14*^, *Fas2*^*G0336*^, and *β-spectrin*^*FY18*^ were induced by incubating larvae or pupae at 37° for two out of every twelve hours over a period of at least two days. Adult females were dissected at least two days after the last heat shock. Any Gal4-UAS driven flies were kept at 29° for 48 hours before dissection.

### Computational Model

Computational modelling was performed using Python, making use of the open-source SciPy library (https://scipy.org/scipylib/). To demonstrate the nonlinearity of the number of misplaced cells with respect to loss of function, misplaced cells were plotted against amount of Nrg or Fas2 removed; an arbitrary unit of one was assigned to the loss of either *Nrg* or *Fas2*. A simple exponential was used as a preliminary model:

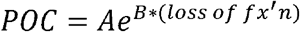

In this model, A and B are arbitrary scale factors. NumPy (https://numpy.org/) was used for array creation and storage. The SciPy optimize.curve_fit tool was used to fit the average and standard deviation of the counts taken for each genotype to our preliminary model. Matplotlib (https://matplotlib.org/), was used to plot the model.

### Immunostaining

Ovaries were fixed for 15 minutes in 10% Paraformaldehyde and 0.2% Tween in Phosphate Buffered Saline. Ovaries were then incubated in 10% Bovine Serum Albumin (in PBS) to block for one hour at room temperature. Primary and secondary immunostainings lasted at least 3 hours in PBS with 0.2% Tween. Three washes (~10 minutes each) in PBS-0.2% Tween were carried out between stainings and after the secondary staining. Primary and secondary antibodies were used at a concentration of 1:150.

### Imaging

Fixed imaging was performed using: a Leica SP5 confocal with a 63x/1.4 HCX PL Apo CS oil lens or an Andor Dragonfly Spinning Disk Confocal microscope with a 60x water objective. Images were collected with LAS AF or the Andor Fusion program respectively. Images were processed (Gaussian blur) using Image J. Minor drift correction was occasionally applied using a custom Python script that included cv2.filter2D, part of the OpenCV library (https://opencv.org/).

### Fluorescence Recovery After Photobleaching

FRAP experiments were performed on a Nikon A1R HD confocal coupled to a Ti2-E inverted microscope. Images were acquired using a Galvano scanner with a 60x/1.49 Apochromat TIRF oil lens using NIS Elements software. Photobleaching of Nrg::YFP was performed on user-defined 0.5 μm^2^ region of interest on cell-cell junctions longer than 2 μm using the 488nm laser at 2 x imaging power at 4 x scan speed. Post-bleaching acquisition of Nrg::YFP was performed at 20 second intervals for 2 minutes, followed by 40 second intervals for 5 minutes, or until the sample moved out of the plane of focus. One image of the imaging field of view was acquired pre-bleaching.

Fluorescence intensity quantification of Nrg::YFP was performed in FIJI ^51^. Average fluorescence intensity was quantified over the 0.5 μm^2^ bleached region for each frame, with xy drift of the ROI corrected manually. FRAP curves were calculated using a double normalization method to correct for imaging acquisition bleaching, analogous to that presented in^52^. The normalized fluorescence intensity of Nrg::YFP at photobleached junctions at each time point *I*_*normFRAP*_(t) was calculated as follows:

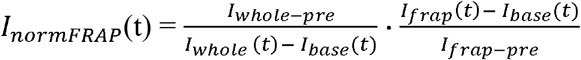

Where

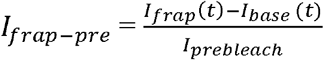

and

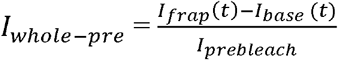

where *I*_*frap*_ the average fluorescence intensity of the photobleached junction region, *I*_*whole*_ is the average fluorescence intensity of the whole field of view (cropped to include only epithelial tissue where appropriate), *I*_*base*_ is the average fluorescence of a ‘background’ region manually defined by drawing a 0.5 μm^2^ region of interest in the cytoplasm of a nearby cell, and *I*_*prebleach*_ is the average fluorescence intensity of the photobleached junction region pre-bleaching. FRAP curves were calculated by calculating a one phase association curve using Prism analysis software (Graphpad).

### Nrg Localization Quantification

Signal intensity was measured along a 3-pixel wide line perpendicular to the center of a cell-cell border. The normalized intensity ratio was calculated as follows:

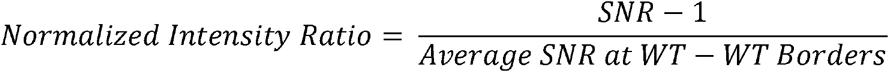

where the signal to noise ratio (SNR) refers to pixel intensity at borders divided by the local background (non-cell-cell border) intensity.

### Statistical analyses

An unpaired, two-tailed students t-test with Welch’s correction was used to determine significance when comparing the number of misplaced cells in an egg chamber. No statistical method was used to predetermine sample size, the experiments were not randomized, and the investigators were not blinded to allocation during experiments and outcome assessment. An unpaired, two-tailed students t-test with Welch’s correction was also used to compare Nrg localization to cell cell borders. The average Normalized Intensity Ratio for each egg chamber was calculated and the distributions of average fold changes were tested for significance. All statistical tests were carried out using GraphPad Prism.

### Misplaced cell counting

Quantification of extralayer cells was performed on Stage 6-8 egg chambers, using at least 3 dissections of at least 5 flies each. For analyses of clonal mutants, the number of extralayer cells was quantified in egg chambers that were at least 60% mutant.

### Spindle Orientation Measurements

Spindle angle determination was performed as previously described ^53^.

### Sequence Alignments

Protein sequence alignments were performed using T-Coffee (http://tcoffee.crg.cat/).

### Reproducibility of Experiments

The *n* for each experiment is provided in Supplemental Table 1.

## Author Contributions

C. Cammarota, T. Finegan, T. Wilson, S. Yang, and D. Bergstralh performed the experiments. C. Cammarota, T. Finegan, and D. Bergstralh designed the experiments. T. Finegan and D. Bergstralh wrote the paper. D. Bergstralh conceived the project.

## Acknowledgements

We are grateful to Holly Lovegrove, Colleen Maillie, Michael Welte and the Welte lab, other members of the Rochester Invertebrate Group, and to Nicole Dawney and other Bergstralh lab members for comments and discussion. We thank Kathryn Neville, Philip Bellomio, Naz Ünsal, and Muskaan Vasandani for technical assistance.

