## Supplemental Information for "An Axon-Pathfinding Mechanism Preserves Epithelial Tissue Integrity"

### Supplemental Figure 1 - Cammarota and Finegan *et al.*

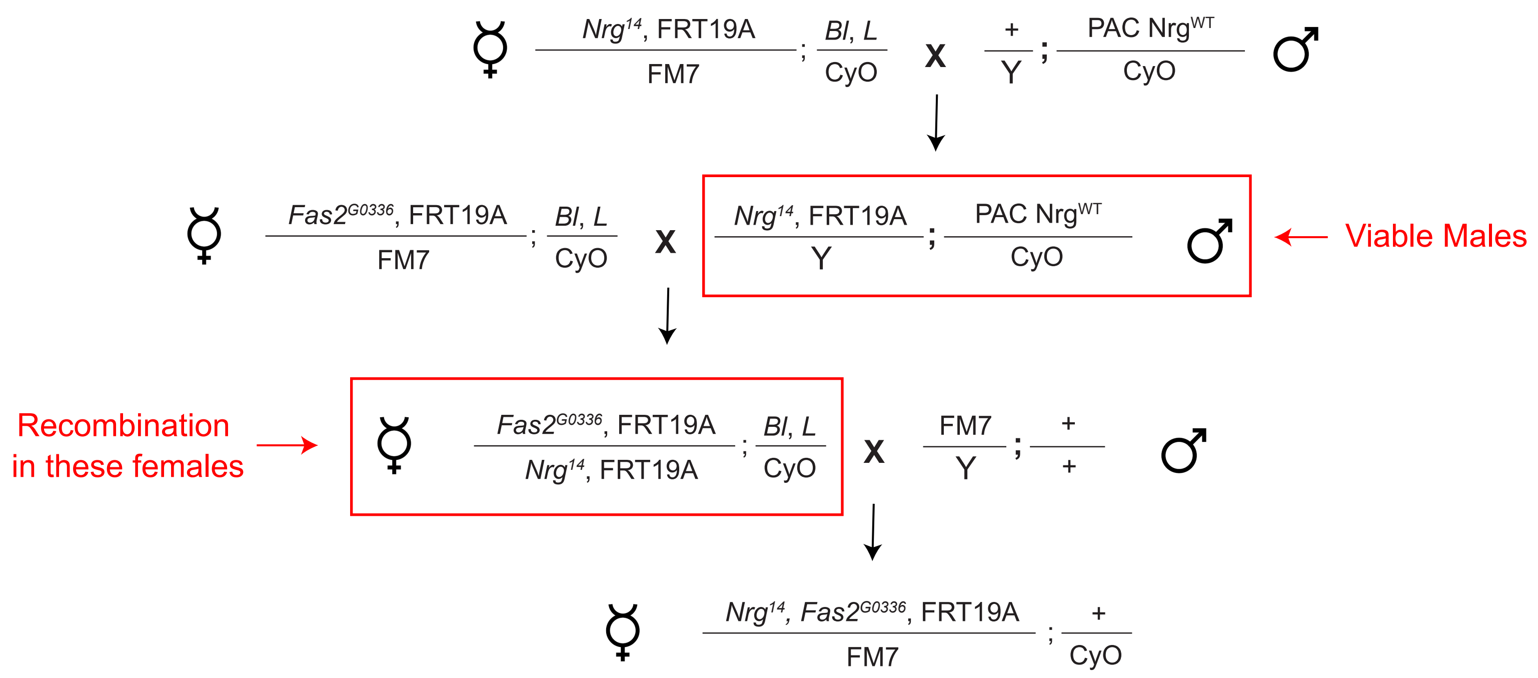

### Supplemental Figure 2 - Cammarota and Finegan *et al.*

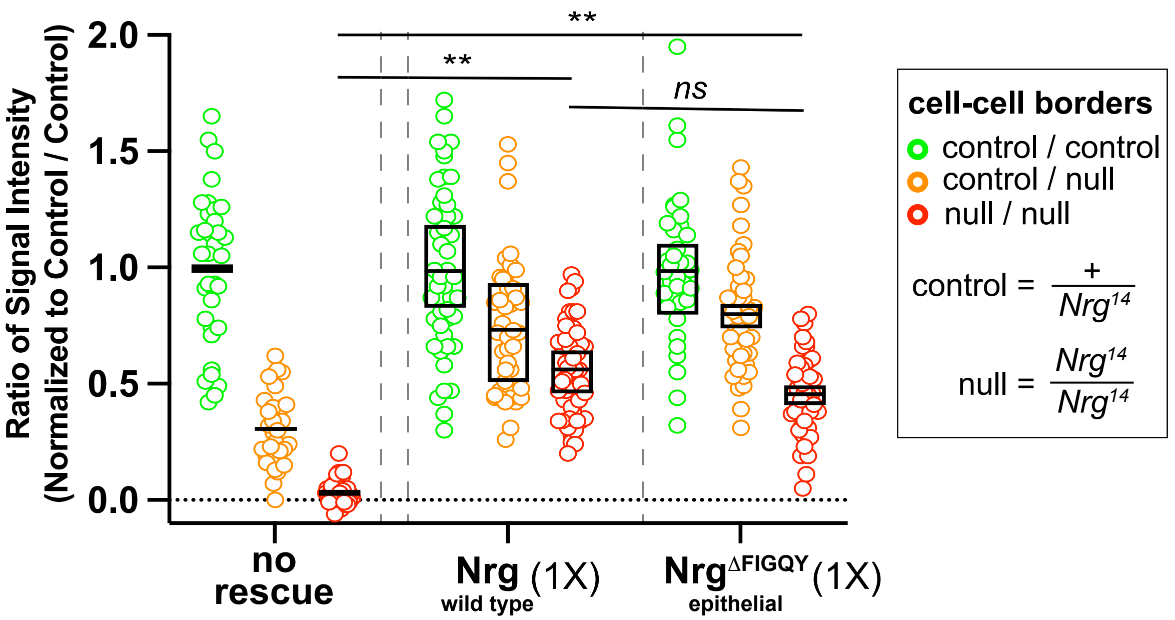

### Supplemental Figure 3 - Cammarota and Finegan *et al.*

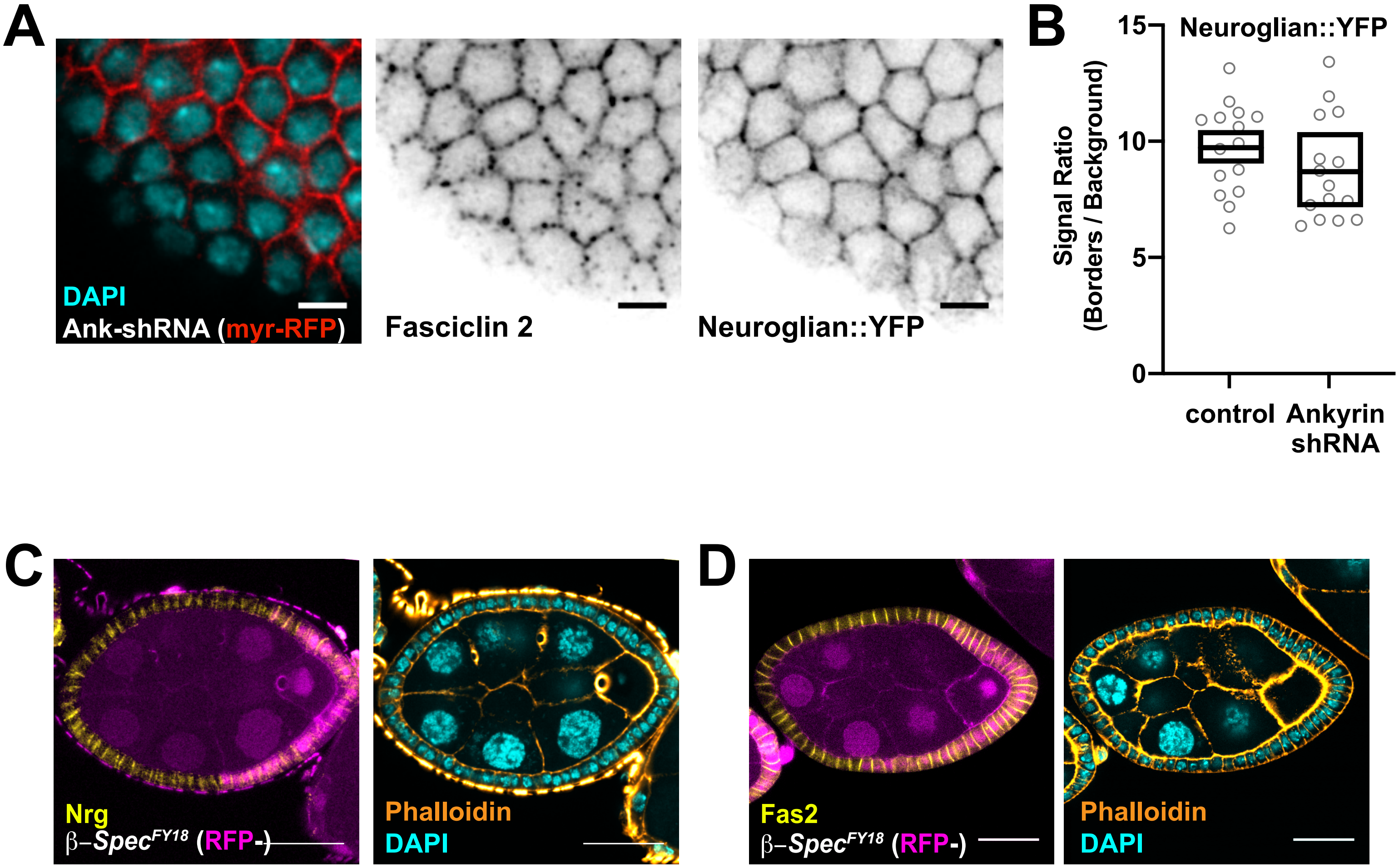

### Supplemental Figure 4 - Cammarota and Finegan *et al.*

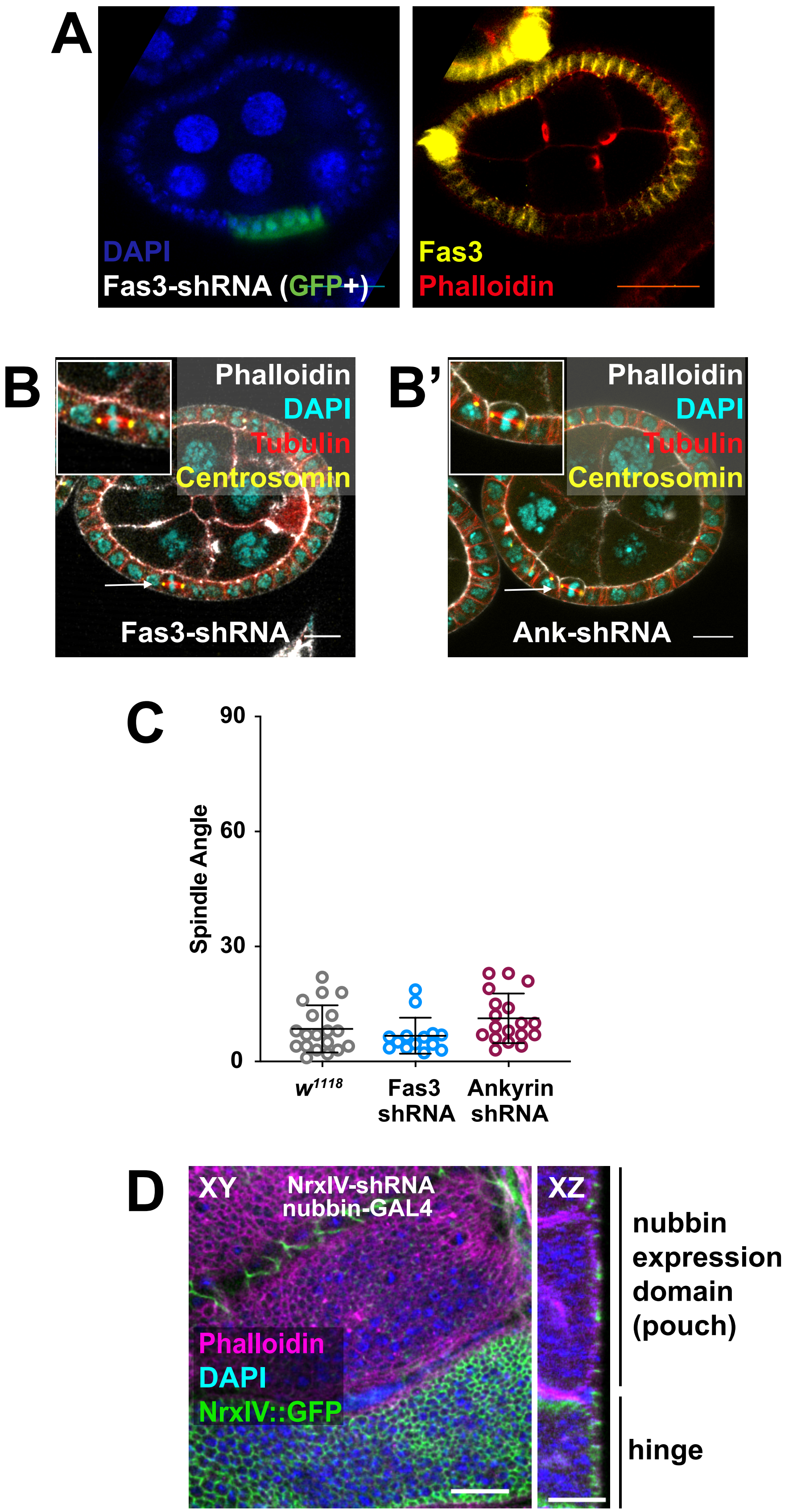

**Supplemental Figure Legends**

**Supplemental Figure 1** – Crossing scheme for the generation of a *Nrg*$^{14}$*, Fas2*$^{G0336}$ double mutant chromosome. Expression of full length Nrg from a genomic rescue construct (P1-derived artificial chromosome $Nrg^{WT}$) on the second chromosome allowed for fertile, viable male flies carrying the lethal null allele *Nrg*$^{14}$.

**Supplemental Figure 2 –** Quantification of Nrg (signal intensity relative to local background) in null and rescue clones. Nrg was visualized by immunostaining. In all conditions, signal intensity correlates with gene dosage; intensity is reduced by ~100% at a null/null border (red, left), but only ~50% when either one copy of $Nrg^{WT}$ or $Nrg^{\Delta FIGQY}$ (epithelial) is expressed in all cells (orange, middle and right). Importantly, total gene dosage in the unrescued condition (1:1, 1:0, 0:0) is not the same in the rescue conditions (2:2, 2:1, 1:1). Signal intensity was normalized against the mean intensity of the control/control condition (green). Therefore, comparisons between genotypes reflect ratios rather than absolute pixel intensity. Statistical tests between egg chambers were performed using an unpaired two-tailed student's t-test. Circles represent individual cell-cell borders. Boxes represent minimum, maximum, and average intensities at the egg chamber level.

**Supplemental Figure 3 –** Fas2 and Nrg localizations are not affected by the loss of Ankyrin or β-spectrin. **A)** Fas2 and Nrg::YFP localization and expression are unchanged by knockdown of Ankyrin. Mosaic expression of UAS-Ankyrin-shRNA and UAS-myrisotylated-RFP (as a marker) were driven in the FE by GR1-GAL4. Fas2 was visualized with the 1D4 antibody. **B)** Quantification of Nrg::YFP signal (relative to local background) at cell borders in egg chambers expressing Ankyrin-shRNA driven by Traffic Jam-GAL4 or egg chambers with the driver alone (control). Three cell-cell borders were measured per egg chamber. Circles represent individual cell-cell borders. Boxes represent minimum, maximum, and average intensities at the egg chamber level. **C and D)** Nrg and Fas2 localization and expression, as measured by immunostaining, is unchanged in *β-Spec*$^{FY18}$ mutant follicle cell clones.

**Supplemental Figure 4** – Verification of Fas3 and NrxIV knockdown and test for spindle orientation effects. **A)** Knockdown of Fas3 expression by shRNA reduces anti-Fas3 immunoreactivity. Fas3 localizes to cell-cell borders during the proliferative stages of follicle cell development. Fas3 is not observed in cells expressing Fas3-shRNA (GFP+). In this experiment, both UAS-GFP and UAS-Fas3-shRNA are driven by Actin-GAL4, which is activated by mitotic recombination-based removal of a stop codon (Flp-out). **B)** Neither disruption of Ankyrin (B) nor Fas3 (B') affects mitotic spindle angle in the follicular epithelium. **C)** Quantification of spindle angles. Mitotic spindles have a mean angle of ~10° relative to the apical surface. **D)** Knockdown of NrxIV expression by shRNA reduces protein expression as measured by fluorescent signal intensity. NrxIV-shRNA was expressed under control of nubbin-Gal4 (expression domain indicated) in the imaginal wing discs of NrxIV::GFP larvae.

#### Supplemental Materials and Methods

##### Fly stocks

The following *Drosophila melanogaster* mutant alleles were used in this study: $Nrg^{14}$ (Eeken et al. 1985), $Fas2^{G0336}$ (Peter et al. 2002), and $\beta\text{-}spectrin^{FY18}$ (Wong et al. 2015). RFP-nls, hsflp, FRT19A was used to make mitotic clones. Recombineered lines expressing $\mathrm{Nrg}^{\mathrm{WT}}$, $\mathrm{Nrg}^{180\Delta\mathrm{FIGQY}}$, and $\mathrm{Nrg}^{167\Delta\mathrm{FIGQY}}$ have been previously described (Enneking et al. 2013). Ectopic protein expression was accomplished using the UAS-GAL4 system (Brand and Perrimon 1993). UAS-Inscuteable (Kraut et al. 1996), UAS-myristoylated-RFP (Andersen et al. 2005), UAS-Fas2-YFP, UAS-Fas2-Extracellular-YFP and UAS-Fas2-Intracellular-YFP (Kohsaka, Takasu, and Nose 2007) have been previously described. We thank the Transgenic RNAi Project at Harvard Medical School (NIH/NIGMS R01-GM084947) for UAS-βHeavy-spectrin shRNA (HMS00882), UAS-NrxIV-shRNA (JF03142), and UAS-Fas3-shRNA (HMC06528) (Perkins et al. 2015). UAS-Ank-shRNA (GD10431) is from the Vienna Stock Center (Dietzl et al. 2007). Expression was driven by Traffic Jam-GAL4 (Olivieri et al. 2010), GR1-GAL4 (Tran and Berg 2003), nubbin-GAL4 (Calleja et al. 1996), or actin5c-FLPout-GAL4 (inducible by FRT/FLP-mediated removal of a stop codon). Neuroglian::YFP (Lowe et al. 2014), and Nrx-IV::eGFP (Buszczak et al. 2007) are Cambridge Protein Trap Insertion and Carnegie Protein Trap Library lines respectively.

**Reagents**

The following antibodies were used in this study: anti-Fas2 (1D4), and anti-Fas3 (7G10) (Developmental Studies Hybridoma Bank), mouse anti-α-tubulin (F2168) (Sigma), rabbit anti-Inscuteable (Kraut et al. 1996)(gift from Jürgen Knoblich), mouse anti-Neuroglian (Bieber et al. 1989) (gift from Michael Hortsch), rabbit anti-Neurexin IV (Stork et al. 2009) (gift from Christian Klämbt), rabbit anti-Centrosomin (Lucas and Raff 2007) (gift from Jordan Raff). Conjugated secondary antibodies were purchased from Thermo Fisher Scientific. Phalloidin was purchased from Invitrogen and Vectashield with DAPI was purchased from Vector Labs.

**Supplemental Table 1**

Sample numbers for each experiment

Fig 1B- Number of Egg Chambers

| | Control | +Inscuteable |
|---|---|---|
| Control | 25 | 22 |
| *Nrg*$^{14}$ | 17 | 34 |
| *Fas2*$^{G0336}$ | 17 | 19 |
| *Nrg*$^{14}$, *Fas2*$^{G0336}$ | 22 | 11 |

Fig 2A- Number of Egg Chambers

| | Control | Inscuteable |
|---|---|---|
| Control | 25 | 22 |
| *Fas2*$^{G0336}$ | 17 | 19 |
| *Fas2*$^{G0336}$ + Nrg$^{WT}$ | 12 | 15 |
| *Nrg*$^{14}$ | 17 | 34 |
| *Nrg*$^{14}$ + Nrg$^{WT}$ | 11 | 25 |
| *Nrg*$^{14}$ + Nrg$^{180\Delta FIGQY}$ | 7 | 10 |
| *Nrg*$^{14}$ + Nrg$^{167\Delta FIGQY}$ | 10 | 24 |

Fig 2D- Number of Borders

| | WT/WT | WT/Mutant | Mutant/Mutant |
|---|---|---|---|
| *Nrg*$^{14}$ (2 ECs) | 35 | 33 | 40 |
| *Nrg*$^{14}$+Nrg$^{WT}$ (3 ECs) | 47 | 43 | 56 |
| *Nrg*$^{14}$+Nrg$^{167\Delta FIGQY}$ (3 ECs) | 37 | 43 | 43 |

Fig 3B- Number of Egg Chambers

| | Control | Ank shRNA |
|---|---|---|
| Control | 25 | 21 |
| *Nrg*$^{14}$ | 17 | 15 |
| *Fas2*$^{G0336}$ | 17 | 13 |

Fig 3C- Number of Borders

| | |
|---|---|
| Control (5 ECs) | 8 |
| Ankyrin shRNA (7 ECs) | 9 |

Fig 4A- Number of Egg Chambers

| | |
|---|---|
| Control | 25 |
| *Nrg*$^{14}$ | 17 |
| *Nrg*$^{14}$ + Fas2$^{TJGal4}$ | 11 |
| *Fas2*$^{G0336}$ | 17 |
| *Fas2*$^{G0336}$ + Fas2$^{TJGal4}$ | 14 |
| *Fas2*$^{G0336}$ + Fas2Intra$^{TJGal4}$ | 13 |
| *Fas2*$^{G0336}$ + Fas2Extra$^{TJGal4}$ | 16 |

Fig 5B-Number of Egg Chambers

| | Control | Fas3 shRNA |
|---|---|---|
| Control | 25 | 30 |
| *Nrg*$^{14}$ | 17 | 28 |
| *Fas2*$^{G0336}$ | 17 | 39 |

Fig 5E- Number of Borders

| | |
|---|---|
| Follicular Epithelium | 20 |
| Wing Disc | 20 |

Fig 5F- Number of Egg Chambers

| | Control | NrxIV shRNA |
|---|---|---|
| Control | 25 | 15 |
| *Nrg*$^{14}$ | 17 | 8 |
| *Fas2*$^{G0336}$ | 17 | 12 |

Supplemental Fig 3B- Number of Borders

| | |
|---|---|
| Control (5 ECs) | 15 |
| Ankyrin shRNA (5 ECs) | 15 |

Supplemental Fig 4C- Number of Spindles

| | |
|---|---|
| Control | 19 |
| *Nrg*$^{14}$ | 14 |
| *Fas2*$^{G0336}$ | 18 |
